# Using time causal quantifiers to characterize sleep stages

**DOI:** 10.1101/550152

**Authors:** Diego M. Mateos, Jaime Gómez-Ramírez, Osvaldo A. Rosso

## Abstract

Sleep plays substantial role in daily cognitive performance, mood and memory. The study of sleep has attracted the interest of neuroscientists, clinicans and the overall population, with increasing number of adults suffering from insufficient amounts of sleep. Sleep is an activity composed of different stages whose temporal dynamics, cycles and inter dependencies are not fully understood. Healthy body function and personal well being, however, depends on proper unfolding and continuance of the sleep cycles. The characterization of the different sleep stages can be undertaken with the development of biomarkers derived from sleep recording. For this purpose, in this work we analyzed single-channel EEG signals from 106 healthy subjects. The signals were quantified using the permutation vector approach using five different information theoretic measures: i) Shannon’s entropy, ii) MPR statistical complexity, iii) Fisher information, iv) Renyí Min-entropy and v) Lempel-Ziv complexity. The results show that all five information theory-based measures make possible to quantify and classify the underlying dynamics of the different sleep stages. In addition to this, we combine these measures to show that planes containing pairs of measures, such as the plane composed of Lempel-Ziv and Shannon, have a better performance for differentiating sleep states than measures used individually for the same purpose.

## 1 Introduction

We think of sleep as our daily period of rest and recovery from the stresses of everyday life; however, research is revealing that sleep is a dynamic activity during which many processes vital to health and well-being take place. Sleep is an active physiological process, while metabolism generally slows down during sleep, all major organs and regulatory systems continue to function normally. New evidence shows that sleep is essential to help maintain mood, memory and cognitive performance [1–3]. Sleep plays also a pivotal role in the normal function of the endocrine and immune systems ([4]). Furthermore, the incidence of sleep problems in different pathologies is uncontroversial and studies show a connection between sleep duration and health conditions such as obesity, diabetes, hypertension, and depression among others ([5–9]). A common taxonomy of sleep differentiates between two different functioning modes: *rapid eye movement* (REM) sleep and *non-REM* sleep (NREM). REM is defined as an active sleeping period marked by intense brain activity. Brain waves are fast and desynchronized similar to those in the waking state, this is also the sleep stage in which most dreams occur. NREM sleep is characterized by a reduction in physiological activity, the brain waves get slower and have greater amplitude, breathing, and heart rate slows down and blood pressure drops. The NREM phase comprises three stages: N1, N2, and N3. The N1 stage is characterized by perceived drowsiness or transition from being awake to falling asleep observed by slowing down the brain waves and muscle activity. Stage N2 is a period of light sleep during which eye movement stops. Brain waves become slower (Theta waves [4-7 Hz]) with occasional bursts of rapid waves (12-14 Hz) called *sleep spindles*, coupled with spontaneous periods of muscle tone mixed. Lastly, stage N3 is characterized by the presence of Delta (0.5-4 Hz) slow waves, interspersed with smaller, faster waves ([10]). In the N3 stage, sleep is deeper, with no eye movement, decreased muscle activity though muscles keep their ability to operate, resembling a coma state. Usually, sleepers pass through these four stages (REM, N1, N2, and N3) cyclically. A complete sleep cycle takes an average of 90 to 110 minutes, with each one lasting between 5 to 15 minutes. These cycles must be maintained for healthy body function in awake state [11].

Polysomnography studies are performed to investigate problems in sleep cycles. They consist of 12 hours of multi-physiological registers such as electroencephalography (EEG), electrocardiogram (ECG), or electrooculography (EOG) among others. Polysomnography studies are usually performed in medical institutions requiring the patient to stay overnight, becoming an issue for the patient, especially when they are children. The detection of the different sleep cycles is done visually (over the electrophysiological signals tracing) requiring specialized clinicians. Automatic classification of sleep stages is an alternative to manual identification which is time-consuming and potentially biased since it depends on personal expertise. The classification of sleep stages by algorithmic means can reduce costs and provide quantitative and therefore comparable assessments.

The existing algorithms for sleep classification can be broadly divided into two groups: multi-parameter (multi-channel EEG, ECG, etc) and single-channel EEG based analysis. The former tend to have better performance [12], but are costly from the patients perspective, for example, in terms of lack of comfort [13] or the sleep disturbances induced by an excessive number of wire connections during the recording process [14]. The latter is more suitable when having in mind the patients comfort, especially given the large amount of time that sleep studies require. According to the evidence [15], single-channel EEG signal can deliver reliable scoring. However, a proper characterization of sleep stages will require more than a reliable signal.

The quantitative identification of sleep features is a promising avenue to help us understand sleep dynamics. A number of methods have been already put to use, among them, time-domain analysis [16, 17], frequency domain [18, 19], time-frequency domain analysis [20, 21] and non-linear feature extraction [19, 22, 23]. However, the candidate features that can be extracted by mathematical and computational means often lack a clear physiological interpretation. In this embarrassment of the riches, the model parameter identification is more often than not through trial and error.

In this work, we propose to use theoretically sounded measures equipped also with straightforward and meaningful physiological interpretation. Specifically, we borrow information-theoretic measures and we use them as functional quantifiers of the probability distribution defined by the EEG signal. Information theory measures can capture the degree to which the neural system integrates information. Particularly, Bandt and Pompe proposed the use of Shannon entropy over a quantification analysis based on ordinal patterns, called *Permutation Entropy* (PE) [24]. This measure has been applied for investigating EEG signals in different contexts as such as epilepsy [25], coma [26], anesthetics [27], and profusely in sleep [28–30], showing better results than traditional analyses. More recently, additional permutation based quantifiers have been used in sleep studies, including *Permutation Min-entropy* (PEmin) [31], *MPR statistical complexity* (Martín-Plastino-Rosso (MPR) complexity) [32], *Fisher information* (FI) [33] and *Permutation Lempel-Ziv complexity* (PLZC) [34].

The paper is organized as follows: In section 2 we explain the characteristics of the dataset including a brief but self-contained introduction to the information quantifiers used. In section 3 we study the distribution of signal patterns in the different sleep stages. Then, we analyze the signals using five different information measures and compare the results. Finally, we analyze the EEG signal through three different complexity/information vs entropy planes. The discussion of the results, possible applications, and future research are discussed in Section 4.

## 2 Methods

### 2.1 Electrophysiological recordings

For this study we used a single-channel EEG signal from a polysomnography recording belonging to 106 subjects taken from the Physionet databank, *The Sleep-EDF Database [Expanded]* ([35, 36]). The dataset is freely available at ([37]). The study group was made up of 54 men and 52 women, 25-85 years of age at the time of the recordings. This polysomnogram (PSGs) collection with accompanying hypnograms (expert annotations of sleep stages) comes from two studies (detail in [36, 38]). The recordings are whole-night polysomnographic sleep recordings containing EEG (from Fpz–Cz electrode locations). The EEG signals were each sampled at 100 Hz. The sleep stages were classified in the five stages above-mentioned: *Awake closed eyes, REM, N1, N2*, and *N3*. Each patient presented a different number of epochs *Ne* per sleep state, being the average number of epochs for each stage: < *N_Awake_* >= 1972, < *N_REM_* >= 169, < *N*_*N* 1_ >= 93, < *N*_*N*2_ >= 483, < *N*_*N*3_ >= 159. All segments have the same length, *L* = 3000 data points (*t* = 30 seconds of recording), and all the recordings were pre-processed with a band-pass filter Fir1 between 0.5 to 40 Hz.

### 2.2 Brief introduction to the information measures

For the study of dynamical phenomena, it is necessary to have a sequence of measurements related to them. These sequences are usually given in the form of time series which allow extracting information on the underlying physical system under study. A time series (i.e. an EEG recording) can be associated with a probability distribution function (PDF), and use information theory quantifiers to characterize its properties. Next, we will introduce the basic notions of the information measures used in this work.

#### Shannon entropy

Given a time series 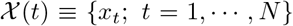, with *N* the number of observations, the Shannon’s logarithmic information measure (*Shannon entropy*) [39] of the associated PDF, *P* ≡ {*p_i_*; *i* = 1, ⋯, *M*} with 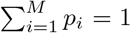, and *M* the number of possible states is defined by:

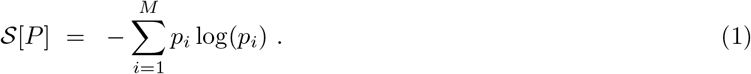

When we have total certainty that our system is in the state *i* the probability *p_i_* = 1 and this functional is equal to zero. In contrast, when the probability distribution is uniform, *P_u_* ≡ {*p_i_* = 1/*M*;∀*i* = 1, ⋯, *M*}, our knowledge about the system is minimum (all the states have the same probability) and the entropy reach its maximum.

#### Rènyi Min-entropy

In 1961, A. Rènyi [40] was looking for the most general definition of information measures, that would preserve the additivity for independent events, and was compatible with the axioms of probability. He defined his generalize entropy as:

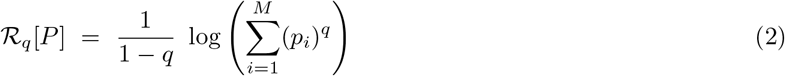

where the order *q* (*q* ≥ 0 and *q* ≠ 1) is a bias parameter: *q* < 1 privileges rare events, while *q* > 1 privileges salient events. The Shannon entropy 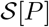 is recovered in the limit as *q* → 1. When *q* → ∞ the Rènyi entropy 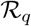 converges to the *Min-entropy* 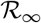

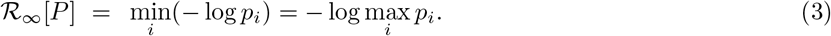

The Min-entropy is the smallest entropy measure in the family of Rènyi entropies. In this sense, it is the strongest way to measure the information content of a discrete random variable. It can be shown that 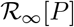 is a function of the highest probability [41]. In particular, Min-entropy is never larger than the Shannon entropy.

#### Fisher Information

The Shannon entropy 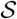 is a measure of “global character”, and in small local regions it lacks sensitivity to strong changes in the PDF. In such a situation, the *Fisher information* 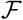 is a more effective quantifier [33, 42]:

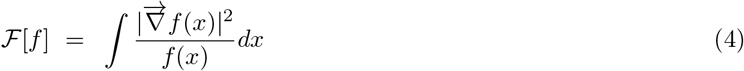

In equation (4), the gradient operator, 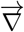, significantly influences the contribution of minute local *f*-variations to the Fisher information value, so that the quantifier is said to be local. Local sensitivity is useful in scenarios whose description necessitates an appeal to a notion of “order” [43–45]. Fisher information constitutes a measure of the gradient content of the distribution *f* (continuous PDF), thus being quite sensitive even to tiny localized perturbations. The intuitive interpretation of the Fisher information is that it measures the ability to estimate a particular parameter and it is also a measure of the state of disorder of a system [42, 46]. Its most important property being the so-called Cramer–Rao bound [47].

For Fisher information measure computation (discrete PDF), we follow the proposal for Dehesa and coworkers in [48] which define 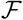 for a discrete distribution as:

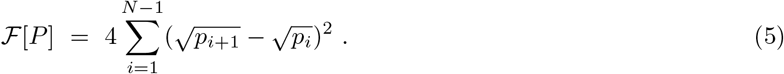

A system in a very ordered state will have a very narrow PDF, the Shannon entropy is 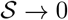 and the Fisher information measure 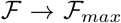. On the other hand, when the system lies in a very disorder distribution (flat PDF) 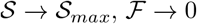. From this simple example we can extrapolate that the Fisher information measure and the Shannon entropy behavior behave reciprocally [49].

#### 2.2.1 MPR Statistical Complexity

If we take the two extremes -perfect order (i.e., a periodic sequence) and a maximal randomness (i.e white noise)- both are very easily to describe (i.e. number of bits [50]) and with complexity closed to zero in both cases. At a given distance from these two extremes, it lies a wide range of possible ordinal structures. Statistical complexity measures [51] allow to quantify this array of behavior [52]. In our work we consider the Martín-Plastino-Rosso (MPR) statistical complexity introduced in [32], because is able to quantify critical details of dynamical processes underlying our data set.

Based in the Lopez-Ruiz et al. work [53], the statistical complexity measure of interest is defined through the functional product form

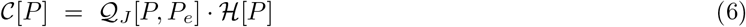

of the normalized Shannon entropy:

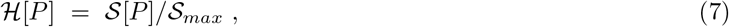

with 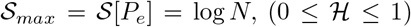 and the disequilibrium 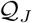 defined in term of the *Jensen–Shannon divergence*. That is,

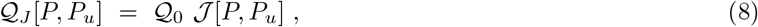

with:

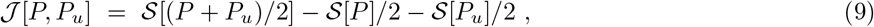

the above-mentioned Jensen–Shannon divergence and 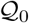, a normalization constant 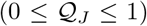, are equal to the inverse of the maximum possible value of 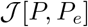. This value is obtained when one of the components of the PDF, *P*, say *p_n_*, is equal to one and the remaining *p_j_* are equal to zero.

The statistical complexity depends on two different probability distributions, the one associated with the system under analysis, *P*, and the uniform distribution, *P_u_*. The distance between this two probability distributions are measured using the Jensen—Shannon divergence [54]. Furthermore, it has been shown that for a given value of 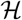, the range of possible 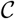 values varies between a minimum 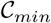 and a maximum 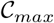, restricting the possible values of the statistical complexity in a given complexity-entropy plane [55]. Because of this, statistical complexity measures can provide important additional information related to the correlational structure between the components of the physical system.

##### Lempel-Ziv complexity

Lempel-Ziv complexity (LZC) is a different way to analyze a sequence; in this case, it is not based on the time series 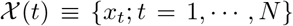 PDF, but in the way that *x_t_* behaves along the sequence. LZC is based on Kolmogorov complexity—the minimal “information” contained in the sequence [47]. To estimate the complexity of 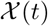 we will use the Lempel and Ziv scheme proposed in 1976 [56]. In this approach, a sequence 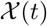 is parsed into a number 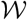 of words by considering any subsequence that has not yet been encountered as a new word. The Lempel–Ziv complexity *c_LZ_* is the minimum number of words 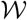 required to reconstruct the information contained in the original time series. For example, the sequence 100110111001010001011 can be parsed in 7 words: 1 · 0 · 01 · 101 · 1100 · 1010 · 001011, giving a complexity *c_LZ_* = 7. An easy way to apply the Lempel-Ziv algorithm can be found in [57]. The LZC can be normalized based in the length N of the discrete sequence and the alphabet length (*α*) as:

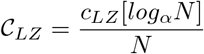

### 2.3 Time series discretization using Bandt-Pompe approach

The study and characterization of time series 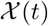 by recourse to information theory tools assume that the underlying PDF is given a priori. A similar problem arises in the Lempel-Ziv complexity context, where it is necessary to have a discrete time series. In the literature there are many methods to quantify continuous time series such as binarization, histograms or wavelet, among others. However, an effective method that emerges naturally is the one introduced by Bandt and Pompe in 2002 called permutation vectors [24].

The permutation vectors method is based on the relative values of the neighbors belonging to the series, and in consequence takes into account the time structure or causality of the process that generated the sequence. To understand this idea, let us consider a real-valued discrete-time series 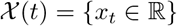, and let *D* ≥ 2 and *τ* ≥ 1 be two integers. They will be called the embedding dimension and the time delay, respectively. From the original time series, we introduce a *D*-dimensional vector 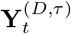:

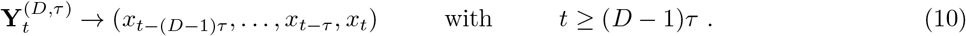

The vector 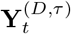 preserves the dynamical properties of the full dynamical system depending on the order conditions specified by *D* and *τ*. The components of the phase space trajectory 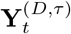 are sorted in ascending order. Then, we can define a *permutation vector*, 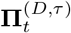, with components given by the original position of the sorted values in ascending order. Each one of these vectors represents a pattern (or motif) with *D*! possible patterns. To clarify, let us show how all this works with an example. Suppose we have a continuous series such as 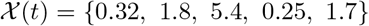 and take the parameters *D* = 3 and *τ* =1. The embedding vectors 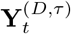 are in this case defined as 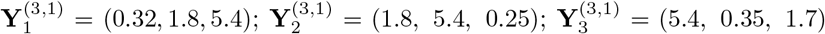, and the respective permutation vectors are 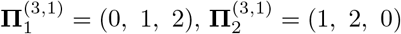 and 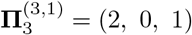.

Regarding the selection of the parameters, Bandt and Pompe [24] suggested working with 3 ≤ *D* ≤ 6 and specifically consider an embedding delay *τ* = 1. Nevertheless, other values of *τ* could provide additional information. It has been recently shown that this parameter is strongly related, if it is relevant, to the intrinsic time scales of the system under analysis [58–60]. In this work we use *D* = 4 and *τ* = 1 for all the analysis, however similar results were obtained for *D* = 5 and *τ* = 1.

The Bandt and Pompe approach applied to information quantifier where used in many works in the past. For each information measures described above, we have their counterpart based on permutation vectors quantification, obtaining: i) Permutation entropy [24], ii) Permutation Min-entropy [58], iii) MPR statistical complexity [61], iv) Permutation Fisher information [45] and v) Permutation Lempel-Ziv complexity [34].

To understand how information quantifiers are applied to permutation vectors it is necessary to generate the rank vectors PDF and then apply each of the different measurements. However, with Lempel-Ziv due to it is an algorithmic measure, it may be a little more difficult to understand. We present an example to clarify this point. Suppose we have a continuous series 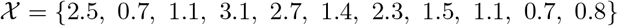 and we use the BP parameter *D* = 2 and *τ* =1, the embedding vectors are 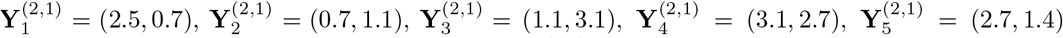, 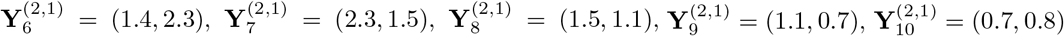. Since *D* = 2, the alphabet has a length equal to *D*! = 2, if the permutation vectors **Π**_0_ = (0, 1) is represented by 0 and **Π**_1_ = (1, 0) by 1 our continuous sequence is quantified as 1001101110, applying the algorithm explain in 2.2.1 and parsing it we get 1 · 0 · 01 · 101 · 101, obtaining 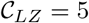. For simplicity, we used *D* = 2 in the example, when *D* > 2 the length of the sequence must be much greater than *D*!. However, as shown in [34] this method can be extended to any *D*.

## 3 Results

### 3.1 Rank vector Analysis

We analyzed 106 EEG signals from 12-hour polysomnography taken from the Physionet databank. The signals were cut in sub-signals correspond to the different periods of sleep stages (*Awake, REM, N1, N2* and *N3*). Figure 1 shows an example of an EEG raw signal (above) and the power spectrum (below) for one typical subject (subject 1) in different sleep stages. All the signals were quantified using the Bandt and Pomp approach with the parameters *D* = 4, 5 and *τ* = 1. These embedding dimension is enough to the capture causality information of the ordinal structure of the time series [24].

**Figure 1:**
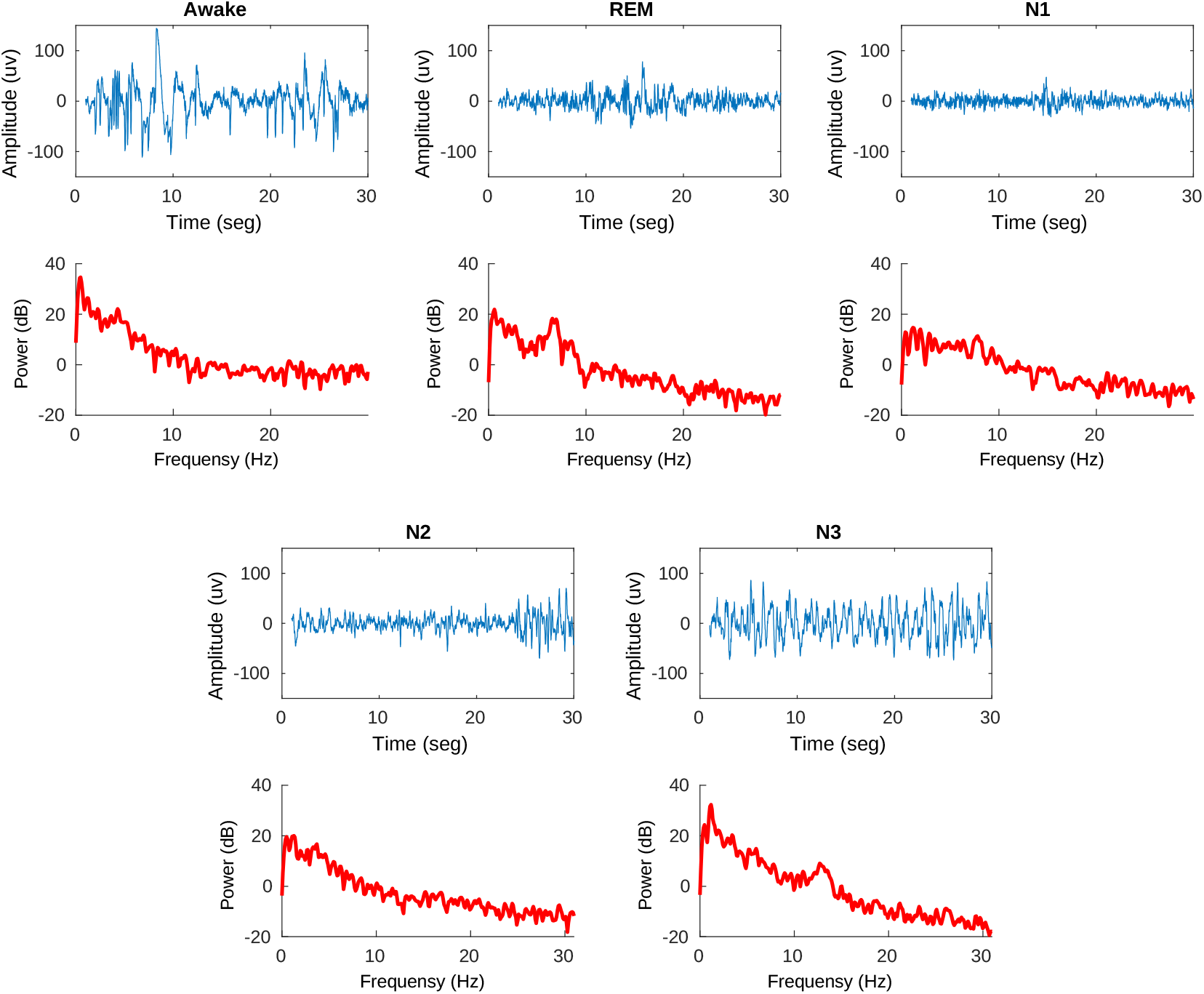
Example of the EEG signals and their respective power spectrum for each sleep stage: Awake, REM, N1, N2, N3.

First, we studied the permutation vectors (**Π**^(*D,τ*)^) distribution in each sleep stage. Figure 2 shows the permutation patterns distribution corresponding to one subject (subject 1), the bars represent the motif average overall epoch in the same stage. Figure 3 shows the same analysis, but the average of the total of 106 subjects. The results in both cases are similar. For subjects in an awake state, the pattern distribution tends to be uniform, the occurrence frequency for the different motifs is comparable for all the cases. In N1, specific patterns start to be more frequent, increasing where the patient enters a deeper sleep state N2. For N3, two particular patterns arise (#1 and #24), they have a higher frequency than the rest which are principally close to zero. This behavior is given because, in an awake state, EEG signals have many superimposed frequencies, being similar to white noises [29]. This results in the necessity of using many different motifs to map the signal, tending the PDF patterns to be uniform. As the person falls asleep, the brain states arise low-frequency waves persistent in time. Particularly, in N1 alpha waves (8-12 Hz) decreases, increasing theta waves (4-7.5 Hz) and starting to appear Low Voltage Mixed Frequency (LVMF) waves. In N2 theta waves are more prominent in time than N1. N3, the deepest sleep stage, presenting over 50 percent delta waves (0-4 Hz) activity [62]. This tendency to exhibit lower frequencies results in the loss of PDF uniformity because we need fewer specific patterns to map the signal. This can be seen more clear in N3 where the predominant patterns are #1 and #24, which correspond to the ascending 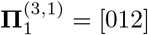 and descending 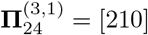 motif. The REM state has a similar PDF to N1 because REM presents theta waves -low-amplitude and mixed-frequency EEG activity-not much different to the waves found in N1 and N2 stages.

**Figure 2:**
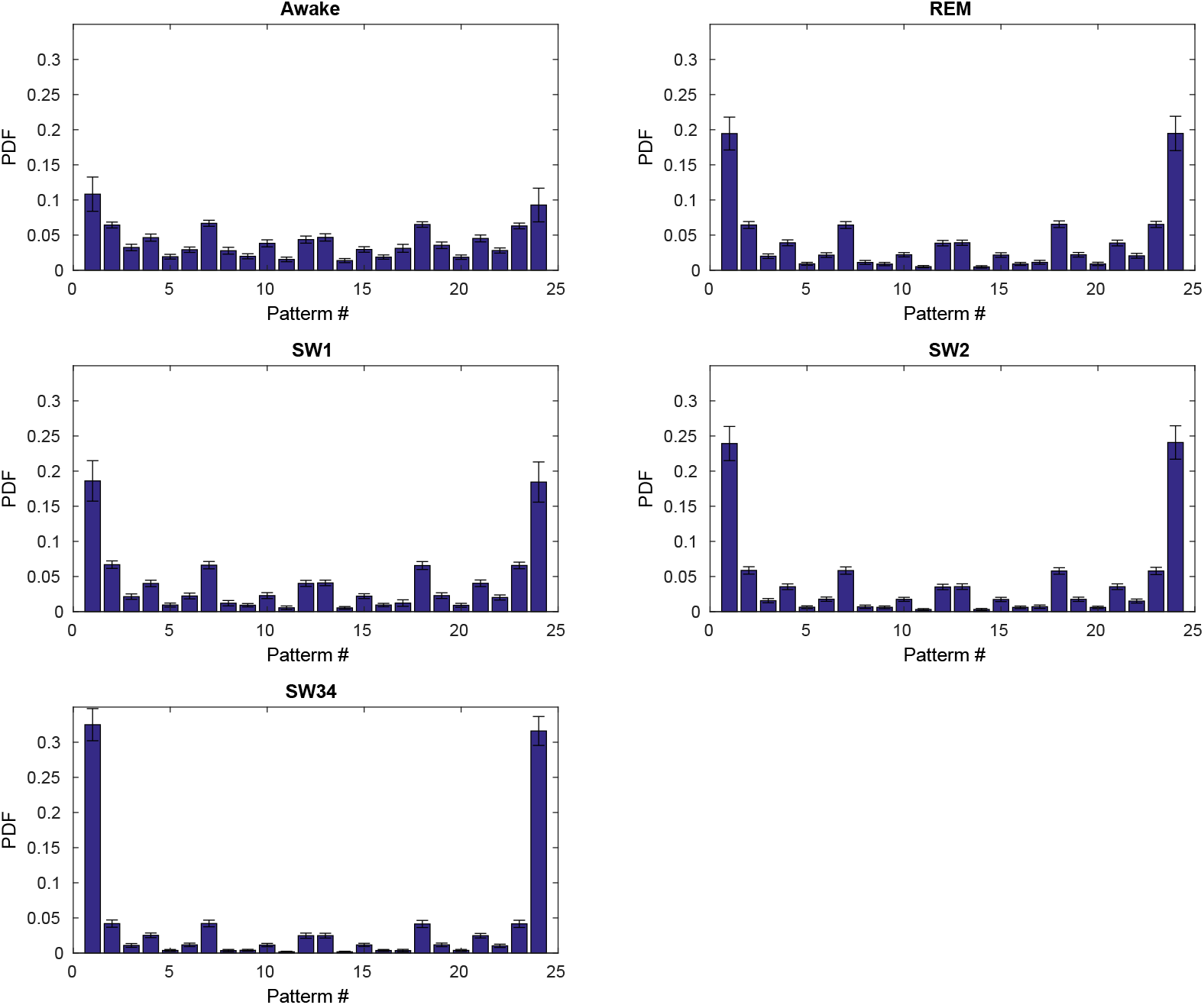
Average Bandt-Pompe PDF for the different sleep states corresponding to subject 1, for *D* = 4 (pattern length) and *τ* = 1 (time lag). The pattern histogram shows the mean value and standard deviation calculated over all segment. Similar results were found for *D* = 5 and *τ* = 1.

**Figure 3:**
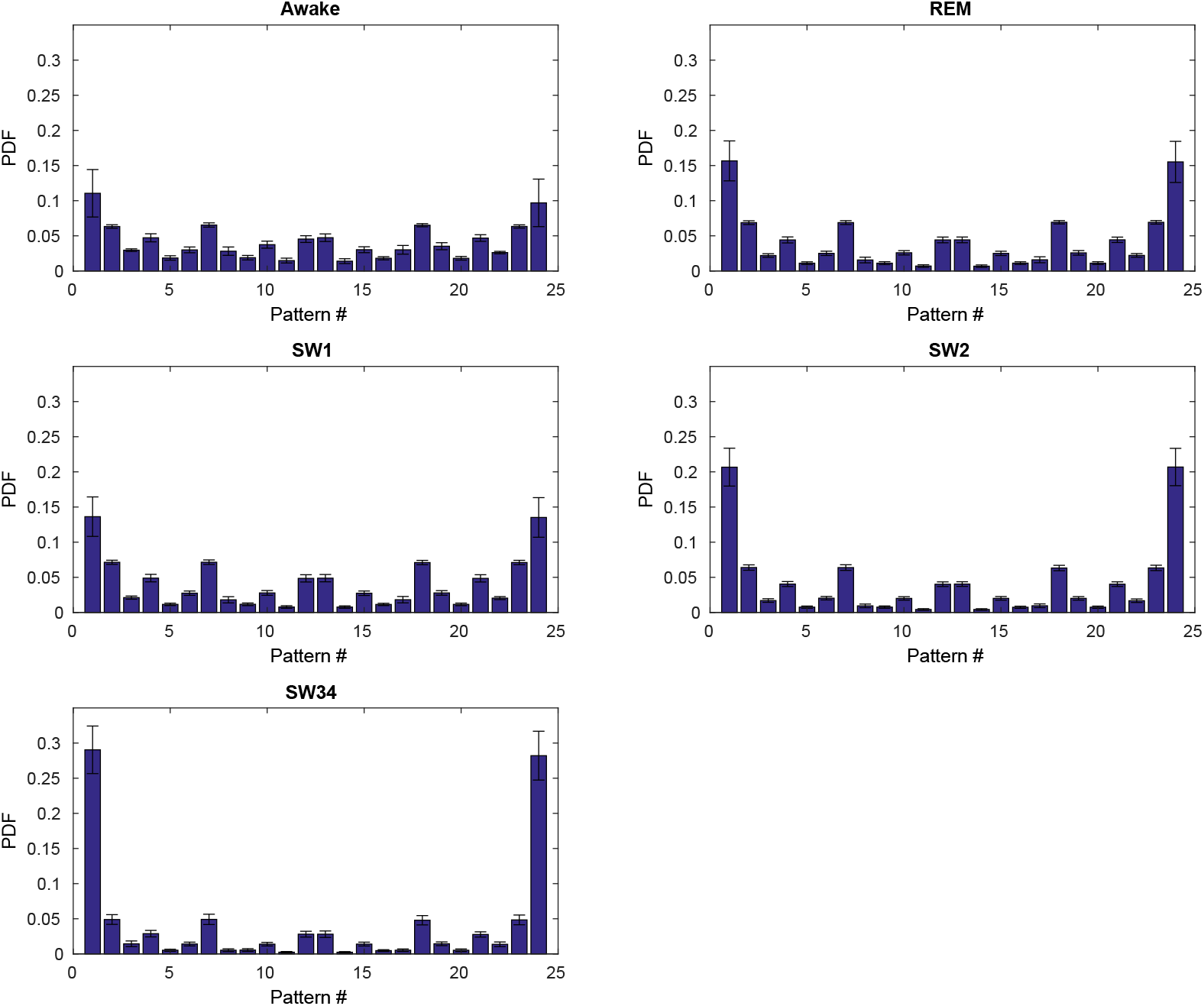
Average Bandt-Pompe PDF for the different sleep states, *D* = 4 (pattern length) and *τ* =1 (time lag). The BP pattern histogram represents the mean value and standard deviation calculated over all subject and all segments. Similar results were found for *D* = 5 and *τ* =1.

### 3.2 Information quantifiers analysis

The following analysis consists of the application of the five measures introduced in section 2 over the PDF rank vectors. Figure 4 depicts a boxplot for A) MPR statistical complexity 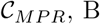 B) permutation entropy 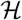, C) Fisher information 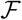, D) permutation Lempel-Ziv Complexity 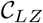 and E) permutation Min-entropy 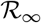. In table 1 we present the quantifiers average values over all patients. The five measures show a remarkable distinction between the different sleep stages. 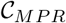 has lower values for subjects in an awake state, increasing as the subjects enter a deeper sleep state. We obtain similar results for 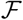 but the values corresponding to the awake state present less dispersion. On the contrary, 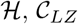 and 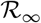, show higher values for awake state, decreasing in REM, N1 and N2 being a minimum for N3. 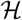 and 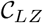 present less value dispersion in comparison with the other quantifiers. Table 2 shows the *p*-value between two state for all quantifiers. All the measures can distinguish significantly between every two states, except for the EEG-N1 case.

**Figure 4:**
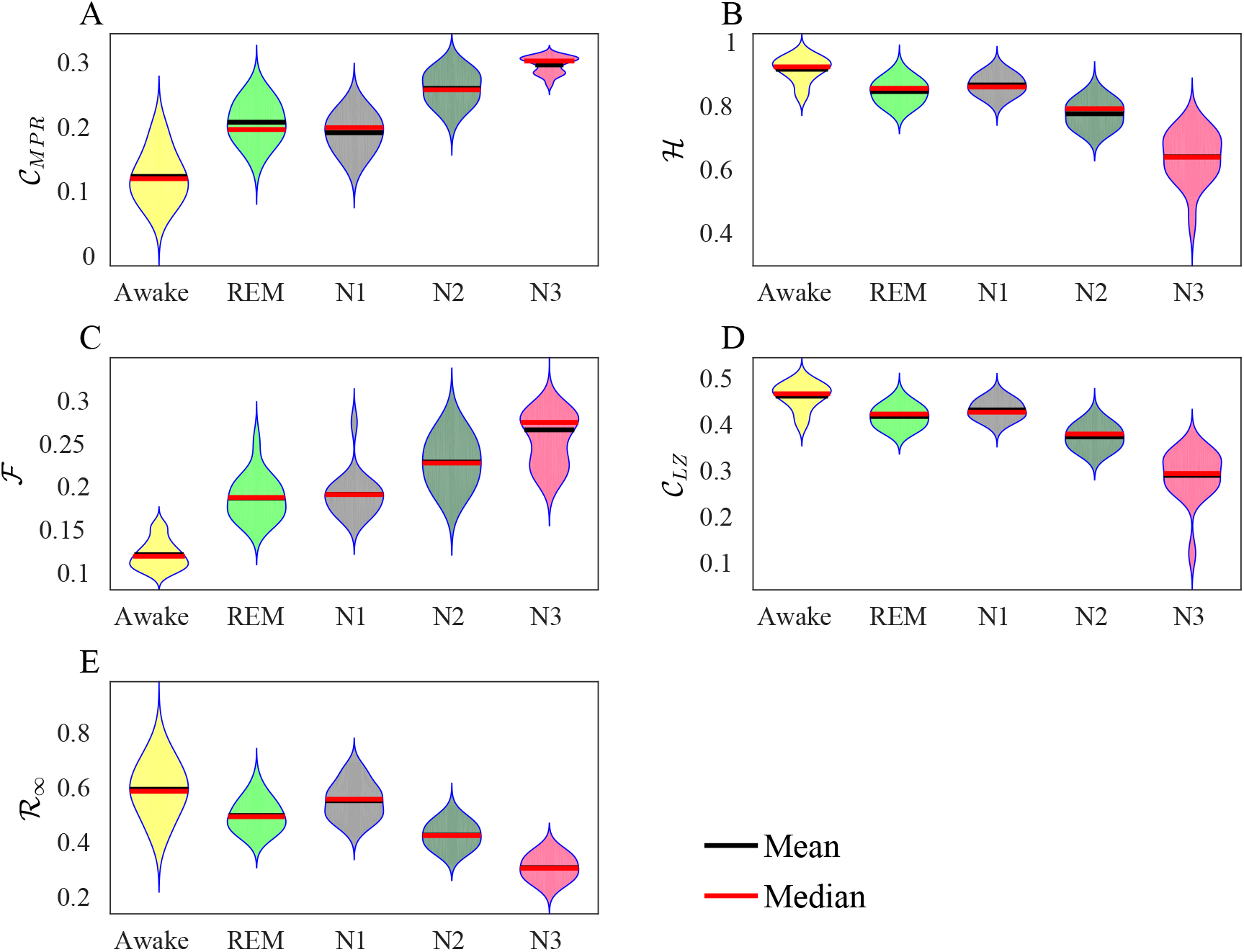
Information quantifiers for different sleep Stages. A) MPR statistical complexity 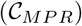, B) normalized Shannon entropy 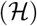, C) Fisher information 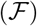, D) permutation Lempel-Ziv complexity 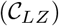, E) permutation Min-Entropy 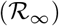. The violin plots were estimated considering the totality of the 106 patients with all their epochs. All the measures used show a clear differentiation between sleep states. The Bandt-Pompe parameter used were *D* = 4 and *τ* = 1, similar results were found using *D* = 5 and *τ* = 1.

**Table 1:** Mean values for all the information quantifiers applied to each sleep state. The quantifiers were calculated over all the segments and all the patients, the parameters were set to *D* = 4 and *τ* = 1.

| Measure | Awake | REM | N1 | N2 | N3 |
| --- | --- | --- | --- | --- | --- |
| $\langle \mathcal{H} \rangle$ | 0.9155 | 0.8453 | 0.8659 | 0.7752 | 0.6395 |
| $\langle \mathcal{C}_{MPR} \rangle$ | 0.1231 | 0.2070 | 0.1908 | 0.2594 | 0.2959 |
| $\langle \mathcal{F} \rangle$ | 0.1209 | 0.1868 | 0.1910 | 0.2286 | 0.2661 |
| $\langle \mathcal{C}_{LZ} \rangle$ | 0.4613 | 0.4173 | 0.4312 | 0.3728 | 0.2893 |
| $\langle \mathcal{R}_\infty \rangle$ | 0.5931 | 0.4973 | 0.5515 | 0.4260 | 0.3077 |

**Table 2:** Information quantifiers *p*-value between two different sleep states. The statistical method used was the Left-tailed Wilcoxon rank sum test. The quantifiers can significantly distinguish the sleep states except for the case REM-N1.

| | | $\mathcal{H}$ | $\mathcal{R}_\infty$ | $\mathcal{C}_{MPR}$ | $\mathcal{F}$ |
| --- | --- | --- | --- | --- | --- |
| Awake | REM | $5.6 \times 10^{-5}$ | 0.003 | $1.9 \times 10^{-5}$ | $7 \times 10^{-7}$ |
| Awake | N1 | $5.7 \times 10^{-4}$ | 0.1 | $3 \times 10^{-5}$ | $7 \times 10^{-7}$ |
| Awake | N2 | $1 \times 10^{-6}$ | $1 \times 10^{-5}$ | $7 \times 10^{-7}$ | $6 \times 10^{-5}$ |
| Awake | N3 | $7 \times 10^{-7}$ | $7 \times 10^{-7}$ | $7 \times 10^{-7}$ | $5.6 \times 10^{-5}$ |
| REM | N1 | 0.148 | 0.011 | 0.168 | 0.637 |
| REM | N2 | $7.4 \times 10^{-5}$ | 0.001 | $6.4 \times 10^{-5}$ | $2 \times 10^{-4}$ |
| REM | N3 | $7.05 \times 10^{-7}$ | $7 \times 10^{-7}$ | $1.9 \times 10^{-7}$ | $3.3 \times 10^{-6}$ |
| N1 | N2 | $2.4 \times 10^{-6}$ | $6.4 \times 10^{-6}$ | $1.9 \times 10^{-6}$ | $3.4 \times 10^{-4}$ |
| N1 | N3 | $7 \times 10^{-7}$ | $7 \times 10^{-7}$ | $1.9 \times 10^{-7}$ | $3.9 \times 10^{-5}$ |
| N2 | N3 | $2.3 \times 10^{-6}$ | $7 \times 10^{-7}$ | $1.9 \times 10^{-5}$ | $8 \times 10^{-3}$ |

In the awake state, we saw that the distribution of the patterns is similar to a uniform PDF (Fig. 4A). This makes the distance between the signal PDF and the uniform PDF minimal, giving low 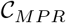 values. A similar result is obtained for the 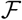, due to no local changes appearing in the PDF of the signal. In the case of 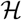 or 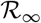, which basically quantify the system’s uncertainty, the values remain maximum because all the states of the signal present equals probabilities. 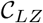 show a marked variability in the occurrence of the permutation patterns resulting in highly complex values due to the impossibility of “compressing” the analysed signal. On the other hand, as the person falls asleep, the uniformity of the PDF rank vectors is lost (see Fig. 4B,C,D). This causes the values of 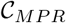 and 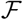 to increase. reaching the maximum in the N3 state. Additionally, the emergence of specific patterns with high probability (especially in N3) in these stages, causes a reduction in the entropic measures (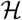 and 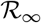). In the same way, when slow waves appear in the N2 and N3 states, the EEG signal violin plot become more monotonous and less complex, giving lower values of 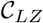.

### 3.3 Causality complexity/information-Entropy maps

Complex/information-entropy maps provide a richer informational repertoire of the system than single quantifiers [44, 61, 63, 64]. In addition, the representation of the results in 2D maps allows a better understanding of the dynamics of the signals. In this work we apply three different complexity/information-entropy maps to the study of EEG signals, 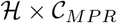 [61], 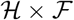 [45] and 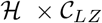 [64].

Figure 5 displays a compilation of the results considered here in the 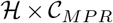 location. The continuous lines represent the curves of maximum and minimum statistical complexity, 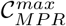 and 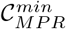, respectively, as functions of the normalized Shannon entropy ([55]). Every dot in the figure represents the mean value distribution for each sleep state. To show entropy-complexity values are not caused by random behaviour of the signal, the recording for all patients were randomized and analyzed. The mean value of the random series was represented in the plane (red star) which lies in the extreme maximum entropy and minimum complexity. Signals that present this kind of behaviour are as different as possible from white noise ([61]). Awake signals have more complexity and less entropy than random signals, but still remain in the plane location where noise signals (poorly structured) lie ([61]). When the sleep state changes to N1, N2 and N3, the complexity tends to increase and the entropy to decrease, moving the values to the centre of the plane.

**Figure 5:**
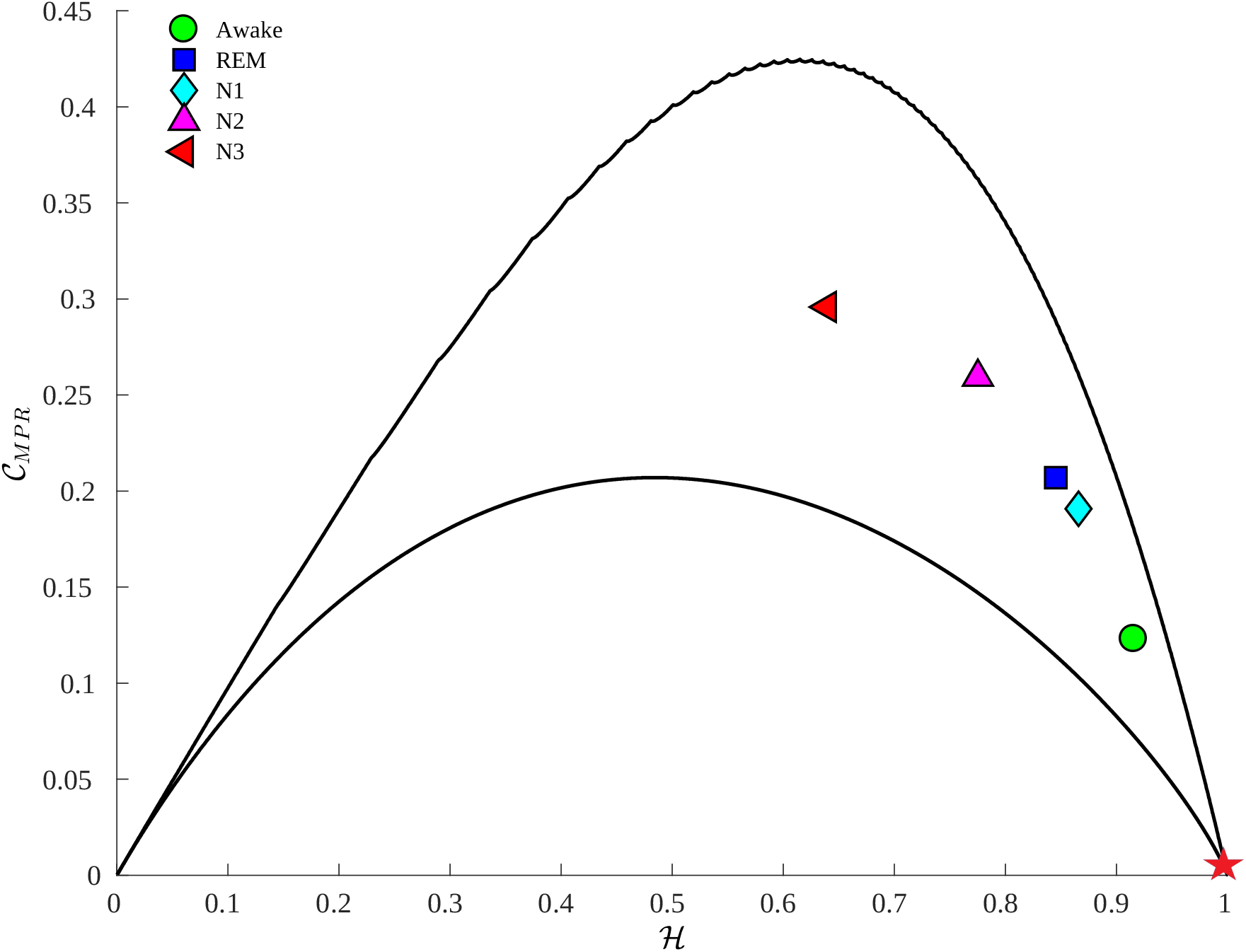
Entropy-Complexity plane 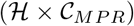 for the 5 different sleep states. Each dot represents the mean value over 106 subjects using the permutation parameters *D* = 4 and *τ* = 1. The awake states are located in a region with greater entropy and less complexity. As the subject falls into deeper sleep stages (N1, N2 and N3) the complexity increases and the entropy decreases. REM is found in intermediate values of complexity and medium-high entropy. All states are clearly discriminated from the randomized signals (red star) which are at the end of the plane. Similar results were obtained for the parameter setting *D* = 5 and *τ* =1.

**Figure 6:**
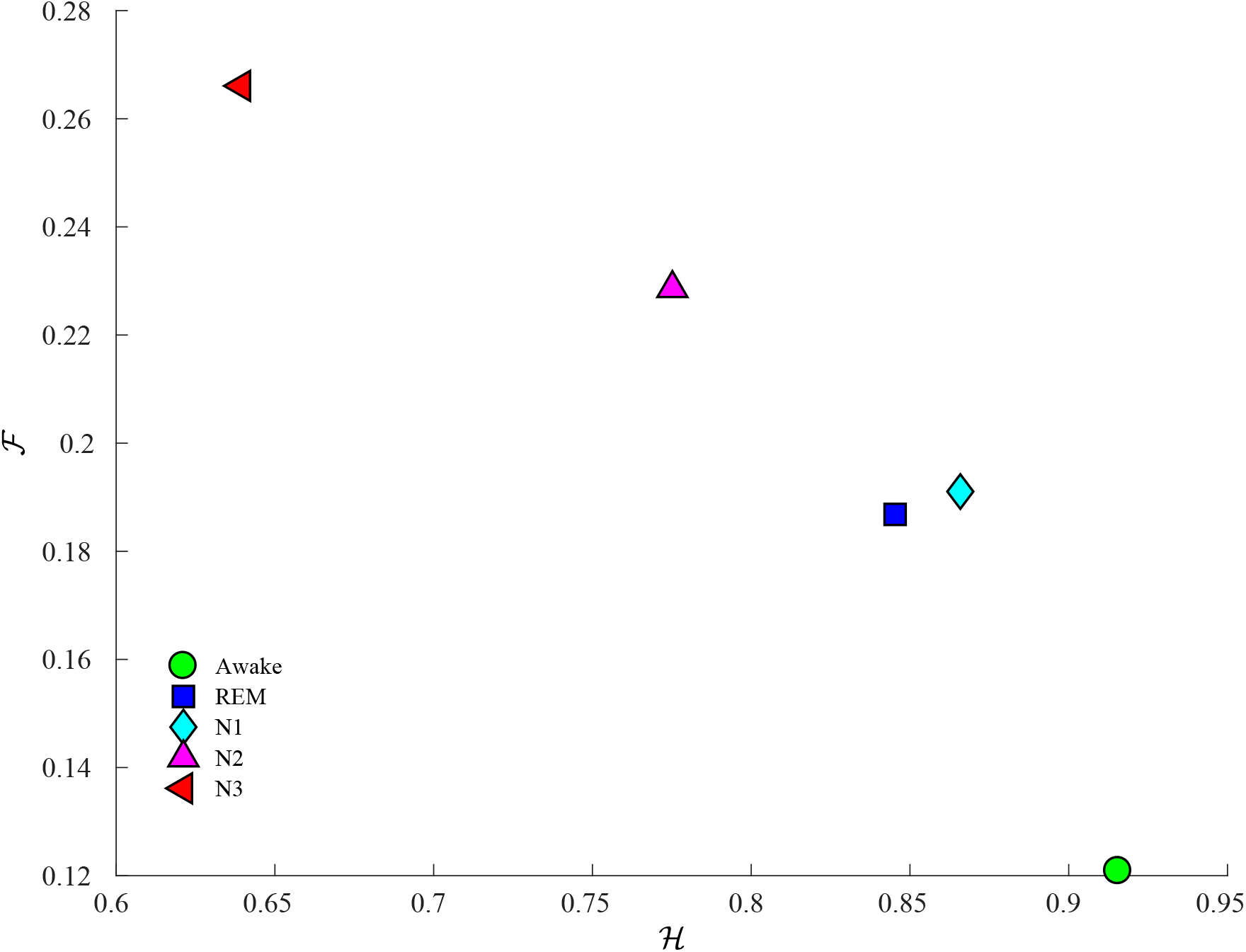
Shannon–Fisher information plane 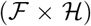 for the 5 different sleep states. Each dot represents the mean value over the 106 subjects using the permutation parameters, *D* = 4 and *τ* = 1. The awake values lies in the high entropy, low information region. REM and sws1 states are located in a region with greater entropy and middle Fisher values. As the subject remains in deeper sleep stages the Fisher values start to increase with the entropy decreasing. Similar results were obtained for the parameters, *D* = 5 and *τ* = 1.

Next, we analyse the signal through the Fisher information - Shannon plane, 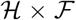. This plane allows to quantify the global versus local characteristic of the time series generated by dynamical processes ([45]). The plane can distinguish between regions corresponding to different sleep states, separating the non-REM from the REM and awake state. However, the REM and N1 states are indistinguishable from each other, especially for Fisher information values. The awake stage shows low 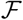 and high 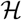. As the sleep becomes deeper, the Non-REM stages are distinguished by having high values of 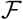 and low values of 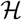.

Finally, we use a plane which combines statistical measures (Shannon entropy) with an algorithmic one (Lempel-Ziv complexity). This 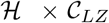 plane has been used to analyse EEG signals in different states of consciousness, giving excellent results characterising different consciousness states such as awake, seizure or coma ([63]). Observing Figure 7, the plane clearly distinguishes all sleep states. The complexity-entropy values live on a straight line over the plane, this is because for random, stationary and ergodic signals–which is the case for EEG under certain conditions ([29])–the Shannon entropy rate tends to the Lempel-Ziv complexity ([56]). The complexity and entropy decrease as the subject falls asleep, being minimal for N3. Unlike the other planes, in this case the values of REM and N1 are clearly differentiable. This renders clear the importance to combine measures for extracting new information about the signal. Similar results were found in intracranial EEG recording ([63]).

**Figure 7:**
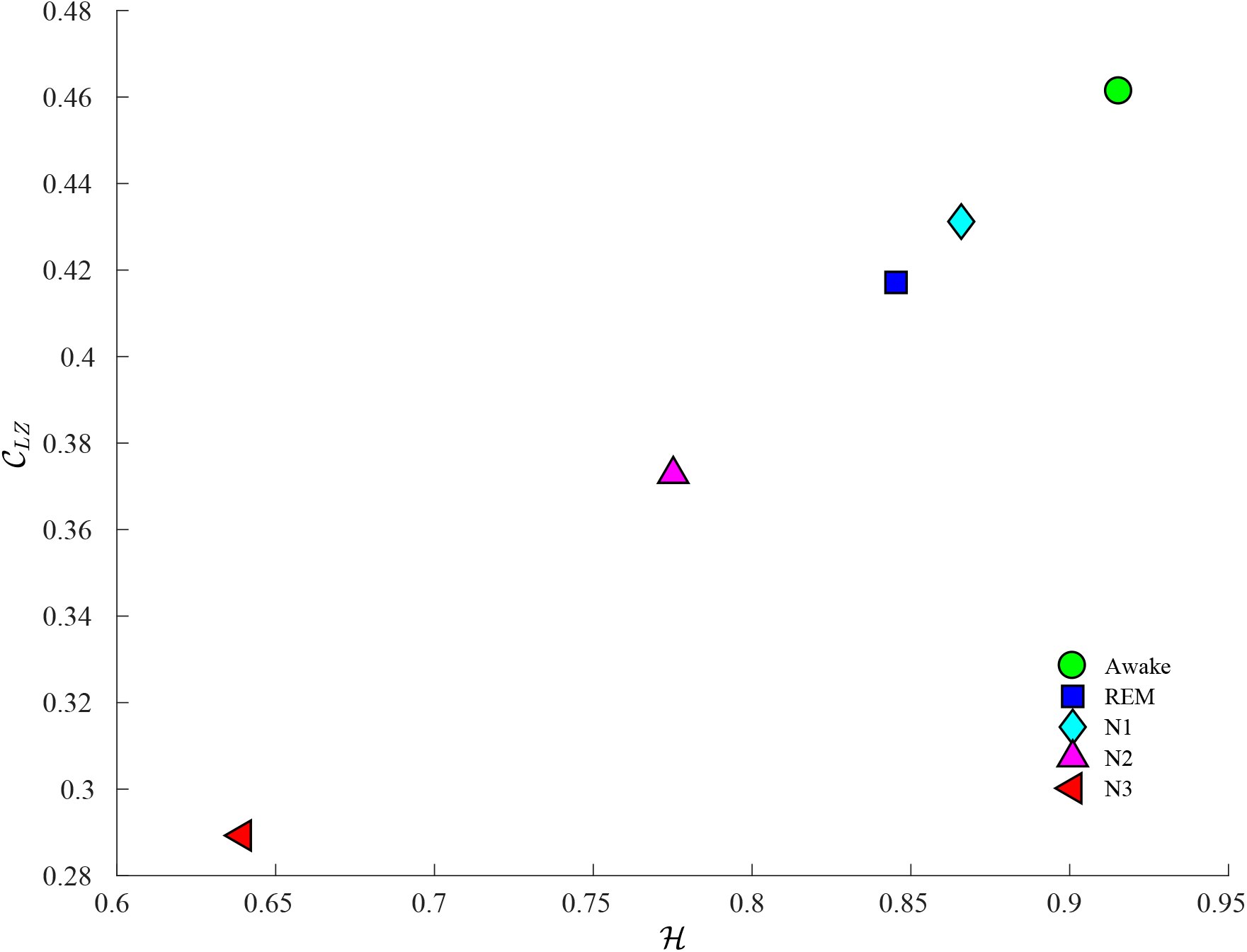
Shannon entropy - Lempel-Ziv complexity plane 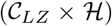 for the 5 different sleep states. Each dot represents the mean value over the 106 subject in the study, using the permutation parameters *D* = 4 and *τ* = 1. The awake stage presents the maximum value of entropy and complexity. As sleep gets deeper, the complexity and entropy decrease been minimum in N3. Similar results were obtained for the parameters, *D* = 5 and *τ* =1.

## 4 Discussion

When the brain is in a wakefulness state, it needs to handle internal and external information, requiring to be able to access to its functions globally. Because of this, the electrical activity becomes more complex with greater variability as observed in the brains waves. This variability in the signal makes that a greater number of motifs are required to map the brain signal, resulting in the permutation vectors PDF converging towards uniformity. This is translated into high values of entropy, both Shannon 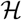 and Renyì 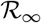. These results are consistent with previous research [28], [65]. MPR complexity shows that signals in the awake state tend to be similar to white noises, corroborating the exposed by Bandt in [29]. Due to the uniformity of the PDF ranks vectors, Fishers information present minimal values. As observed for the awake state, the complexity of Lempel-Ziv is high due to the necessity of a large number of different patterns needed to encode the signal. Similar results were found in previous studies [34, 63].

As sleep progresses, most of the vital signals decrease (respiration, blood pressure, etc). The cortex becomes less active, so the EEG signal becomes slower and more repetitive with a marked predominance of delta waves. Delta waves are slow and more monotonous, making that the number of patterns required to mapping the signal decreased. Some motifs become more important than others, specially those mapping the complete ascent or descent of the signal. This is reflected in a decrease in the signal entropy both 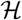 and 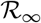. Due to this regularity in the EEG, the code required to reconstruct the signal becomes smaller, producing a drop in the permutation Lempel-Ziv complexity. Similar results were found in previous works in intracranial EEG. This behaviour is not exclusive to the quantification by Rank vectors, similar results have been obtained different quantification techniques [65, 66]. The predominance of slow waves in No-REM states produces a sort of brake in the uniformity PDF giving more probability to specific patterns, which in turn results in an increase in MPR complexity and the Fisher information.

We have seen that the differentiation between REM and N1 status is not an easy task (see 4). This is expected considering that the REM signal contains frequencies that are present in the “awake state and in the lighter stages of sleep N1 [28]. The presence of 11-16 Hz. activity (sleep spindles) in N1, and more abundant alpha activity (8-13Hz) in REM sleep means that these two stages present activity at an overlapping frequency range, which explains the proximity of the information values obtained for these two stages. Difficulty in separating N1 and REM sleep has also been encountered using other measures [67]. Despite the similarities of the REM and N1, there are still enough differences between them. These can be quantified by 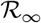 or better through the complexity/information-entropy planes, especially for the 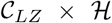 plane shown here.

The use of complexity/information-entropy planes provides a better data visualisation tool, containing information about the signals which are impossible to get applying each measure individually. The three planes presented in this work show a clear differentiation over all sleep states. In particular, they allow us to differentiate N1 from REM. It is thanks to the combination of different approaches that these results could be obtained. For example, i) statistical and algorithmic measures 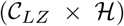, ii) local and global measures 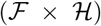 or iii) order and disorder dynamics 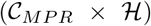.

This work makes a substantiated case for the use of information-theoretic tools for the analysis of the one-channel EEG signals, complementing the visual inspection adopted by electrophysiologists. The information quantifiers used have proven highly effective for the detection and classification of sleep states. Provided the possibility of working with short sequences, its simplicity and efficient calculation make them suitable to use in clinical settings requiring real-time. This feature would allow to develop technical devices of use in sleep studies, without the subject having to check in a hospital or sleep clinic. Given the importance that sleep cycles have in peoples health, in future work we will focus on applying these analytical methods for the study of pathological EEGs. To that we will explore data from patients with insomnia, circadian rhythmic sleep-awake and parasomnia among others, using these quantifiers to validate them as potential biomarkers.

## 5 Conclusion

The study of sleep stages are of vital importance to understand both healthy and pathological conditions. Dysfunctions in sleep cycles generate problems such as fatigue, muscular pains and attention problem among others. Because of that, the detection and proper characterisation of sleep stages is of paramount importance. In this work we have been able to characterise classify the dynamics of different sleep stages in an easy and fast way, using measures borrowed from Information Theory. Out results show that the information-theoretic measures studied here -Shannon’s entropy, MPR statistical complexity, Fisher information, Rènyi Min-entropy and Lempel-Ziv complexity-represent a promising avenue to foster our understanding of sleep cycles in quantitative basis, facilitating the development of low-cost equipment for signal processing applied to human sleep. In particular, we find that the information quantifiers proposed here can be effectively used for the detection and classification of sleep states. Furthermore, the quantifiers are robust to short signal sequences and fast and straightforward to compute which make them suitable to be used in real time analysis of sleep recordings.

